# Motor resonance is modulated by an object’s weight distribution

**DOI:** 10.1101/2020.10.11.335000

**Authors:** Guy Rens, Jean-Jacques Orban de Xivry, Marco Davare, Vonne van Polanen

## Abstract

Transcranial magnetic stimulation (TMS) studies showed that corticospinal excitability (CSE) is modulated during observation of object lifting, an effect termed ‘motor resonance’. Specifically, motor resonance is driven by movement features indicating object weight, such as object size or observed movement kinematics. We investigated in 16 humans (8 females) whether motor resonance is also modulated by an object’s weight distribution. Participants were asked to lift an inverted T-shaped manipulandum with interchangeable center of mass after first observing an actor lift the same manipulandum. Participants and actor were instructed to minimize object roll and rely on constrained digit positioning during lifting. Constrained positioning was either collinear (i.e., fingertips on the same height) or noncollinear (i.e., fingertip on the heavy side higher than the one on the light side). The center of mass changed unpredictably before the actor’s lifts and participants were explained that their weight distribution always matched the actor’s one. Last, TMS was applied during both lift observation and planning of lift actions. Our results showed that CSE was similarly modulated during lift observation and planning: when participants observed or planned lifts in which the weight distribution was asymmetrically right-sided, CSE recorded from the thumb muscles was significantly increased compared to when the weight distribution was left-sided. During both lift observation and planning, this increase seemed to be primarily driven by the weight distribution and not specifically by the (observed) digit positioning or muscle contraction. In conclusion, our results indicate that complex intrinsic object properties such as weight distributions can modulate activation of the motor system during both observation and planning of lifting actions.

**Highlights:**

• Motor resonance is observation-induced activity in the observer’s motor system
• We used a dyadic lifting task of objects with asymmetrical weight distribution
• We investigated which movement features modulate motor resonance
• Motor resonance is modulated by the object’s weight distribution
• Motor resonance is driven by observed and planned digit positioning

## Introduction

Skilled object manipulation not only relies on tactile feedback but also on anticipatory mechanisms (Johansson & Westling, 1988). However, predictive lifting errors are made when object weight is wrongly estimated (e.g., lifting an opaque box with unexpected amount of filling). In this situation, individuals update their ‘sensorimotor memory’ which contains short-term associations between previous hand-object experiences and the visual object properties (Baugh et al., 2012). As such, the sensorimotor memory can be flexibly updated and subsequently used for predicting object weight and planning skilled hand-object interactions.

Importantly, skilled hand-object interactions not only require accurate planning for object weight but also for weight distribution. For instance, when having to avoid content spill, object roll has to be minimized by generating appropriate compensatory torque to offset external torque induced by an unbalanced weight distribution. Lukos et al. (2007) showed that individuals can update their sensorimotor memory for an object’s weight distribution, in turn enabling them to predictively generate appropriate compensatory torque. In addition, Fu et al. (2010) showed that individuals scale their fingertip forces in function of their digit positioning: when digit positioning is constrained, individuals scale their fingertip forces in function of these fixed contact points. Conversely, when digit positioning is unconstrained, individuals scale their fingertip forces accurately in function of their self-chosen contact points. However, it has been argued that, in line with these findings, object lifting with unconstrained digit positioning relies on both predictive and feedback driven mechanisms: although fingertips are positioned based on the initial motor command, trial-to-trial variability in actual positioning is induced by contextual and executional noise. As a result, the initially planned fingertip forces need to be updated in function of tactile feedback about the actual positioning (Mojtahedi et al., 2015). To end, it is interesting to note that Lukos et al. (2013) showed that when individuals plan to lift an object with unpredictable weight distribution, they rather rely on a motor command based on their previous lift (i.e., sensorimotor memory) than on a generic or ‘neutral’ motor command. Combined, these studies show that individuals can plan dexterous manipulation of objects with complex properties but that they do rely on sensorimotor integration based on their previous trial (Lukos et al., 2013) and on real-time haptic feedback during lift execution (Mojtahedi et al., 2015).

Performing hand-object interactions modulates activity within the motor system during both motor execution and planning (for a review see Hannah, 2020). That is, corticospinal excitability (CSE) probed with transcranial magnetic stimulation (TMS) over the primary motor cortex (M1) is modulated during motor tasks. Loh et al. (2010) showed that when individuals plan to lift an object, CSE initially reflects the weight of the previously lifted object. However, when a visual cue indicates that the object weight has changed, CSE modulation is altered and becomes representative of this new weight (Loh et al., 2010). In Davare et al. (2019), participants were instructed to grasp and lift an asymmetrical weight distribution either with constrained or unconstrained digit positioning. They showed that CSE was increased when motor predictability was lower (i.e., unconstrained positioning). Noteworthy, these effects were only present after object contact but not during reaching. As such, their findings suggest that CSE modulation during early contact does not only reflect sensorimotor integration of the previous trial but also on-line feedback about digit placement.

Parikh et al. (2014) showed that when individuals plan to grasp (but not lift) an object, CSE is decreased when planning to exert high compared to low force. In addition, the authors argued that, considering these effects were not altered by paired-pulse TMS, that CSE modulation appears to be driven by inputs from regions outside M1. To investigate the role of the somatosensory cortex (S1) in predictive lift planning, Parikh et al. (2020) asked participants to lift an asymmetrical weight distribution after virtually disrupting either M1 or S1 with repetitive TMS. They found that when digit positioning is constrained, force planning relies on memory retrieval in M1. In addition, when digit positioning is unconstrained, grasp planning relies on memory retrieval of digit positioning in M1 and on integrating haptic feedback regarding digit positioning to generate appropriate load forces in S1. Taken together, these studies show that M1 and S1 are involved in sensorimotor integration during the planning and execution of hand-object interactions (Loh et al., 2010; Parikh et al., 2014) and, in particular, also on objects with an asymmetrical weight distribution (Davare et al., 2019; Parikh et al., 2020).

Although execution of hand-object interactions is pivotal in rapidly updating the sensorimotor memory, other studies have shown that humans are able to generate similar representations during observation of object lifting as well. For instance, Meulenbroek et al. (2007) demonstrated that when two individuals incorrectly predict an object’s weight, the second individual will make a smaller lifting error after observing the first individual making one. These findings have been supported by other studies (Buckingham et al., 2014; Reichelt et al., 2013). In addition, it has been shown that observation of skilled lifting can improve predictive object lifting as well, albeit in a smaller manner than observing lifting errors (Rens et al., 2020, 2021; Rens & Davare, 2019)

Akin to motor execution, observing hand-object interactions not only alters sensorimotor representations but also activates the observer’s motor system (for a review see Naish et al. 2014). Fadiga et al. (1995) were the first to demonstrate that CSE is similarly modulated during the execution and observation of hand actions. They argued that the motor system is potentially involved in action understanding through a bottom-up mapping (‘mirroring’) of observed actions onto the same cortical areas involved in their execution (for a review see: Rizzolatti et al., 2014). Consequently, action observation-driven modulation of CSE has been termed ‘motor resonance’. With specific interest to observation of object lifting, Alaerts et al. (2009, 2010a, 2010b) demonstrated that motor resonance is modulated by observed movement features indicating object weight, such as intrinsic object properties (e.g., size), muscle contractions and movement kinematics. Specifically, CSE is increased when observing lifts of heavy compared to light objects. Critically, other studies have shown that these motor resonance effects are not robust. For instance, Buckingham et al. (2014) demonstrated, using the size-weight illusion, that CSE modulation is driven by object size when observing skilled but not erroneous lifts. In addition, Tidoni et al. (2013) showed that motor resonance is differently modulated when observing an actor with truthful or deceptive intentions. Last, Rens et al. (2020) demonstrated that motor resonance is easily biased by differences within the contextual setting even though the observed lifting actions are the same. In addition, they showed that this bias was generated by top-down inputs from the posterior temporal sulcus.

Although Alaerts et al. (2010) and Buckingham et al. (2014) highlight that motor resonance can reflect simple object properties, it is unknown whether it can reflect more complex ones such as an object’s weight distribution. Due to the importance of accurately estimating an object’s weight (i.e., avoid damage) and weight distribution (i.e., avoid content spill), it is plausible that they rely on similar mechanisms for estimating these properties. However, as minimizing object roll requires a valid digit position-force coordination pattern, the observer’s motor system should integrate both observed digit positioning and forces (rather than solely encoding force) for accurately encoding the weight distribution. As such, in this study we wanted to investigate whether the observer’s motor system encodes (a) observed force scaling, (b) observed digit positioning or (c) the weight distribution, as indicated by a combination of these features. We asked participants to grasp and lift an inverted T-shaped manipulandum with interchangeable (left, middle or right) center of mass after first observing an actor lift the same manipulandum. Participants and actor were required to minimize object roll during lifting by generating appropriate compensatory torque. The center of mass changed unpredictably before the actor trials, but participants were informed that they would always lift the same weight distribution as the actor. As such, participants could potentially estimate the object’s center of mass during lift observation and use this information to predictively plan their own lifts. For lifting the asymmetrical weight distributions, we constrained digit positioning to two distinct possibilities: (1) Placing the fingertips on the same height (‘collinear positioning’) or placing the finger on the heavy side higher than the one on the light side (‘noncollinear positioning’). Importantly, when compensatory torque is generated for an asymmetrical weight distribution, the fingertip on the heavy side generates more force when using a collinear positioning compared to a noncollinear one (Fu et al., 2010). By relying on these constrained digit positionings and the associated force requirements for skilled lifting, we hypothesized that we could disentangle whether the motor system encodes force or digit positioning during lift observation. For doing so, we used TMS to probe CSE during lift observation and planning.

In line with Alaerts et al. (2010), we hypothesized that, if the observer’s motor system encodes force exertion by the fingertips, motor resonance should be significantly increased when observing skilled lifts on the asymmetrical weight distributions with collinear compared to noncollinear positioning. For instance, when the index finger is on the heavy side, it generates more force in the collinear condition than in the noncollinear one. As such, CSE recorded from the index finger should be larger in the collinear compared to noncollinear condition. In contrast, if motor resonance is primarily modulated by digit positioning, we would expect that motor resonance, when observing lifts on the asymmetrical weight distribution, would be increased when observing lifts with noncollinear compared to collinear positioning.

## Methods

### Participants

16 individuals participated in the present study (8 females; mean age = 24 ± 3 years). The Edinburgh Handedness Questionnaire (Oldfield, 1971) revealed that all participants were strongly right-handed (> 90). Prior to participation, participants were required to fill in a TMS safety screen questionnaire based on Rossi et al. (2011). Moreover, all participants had normal or corrected-to-normal vision, were free of neurological disorders and had no motor impairments of the right upper limb. Participants gave written informed consent and were financially compensated for their time. The protocol was in accordance with the Declaration of Helsinki and was approved by the local ethical committee of KU Leuven, Belgium.

**Figure 1.**
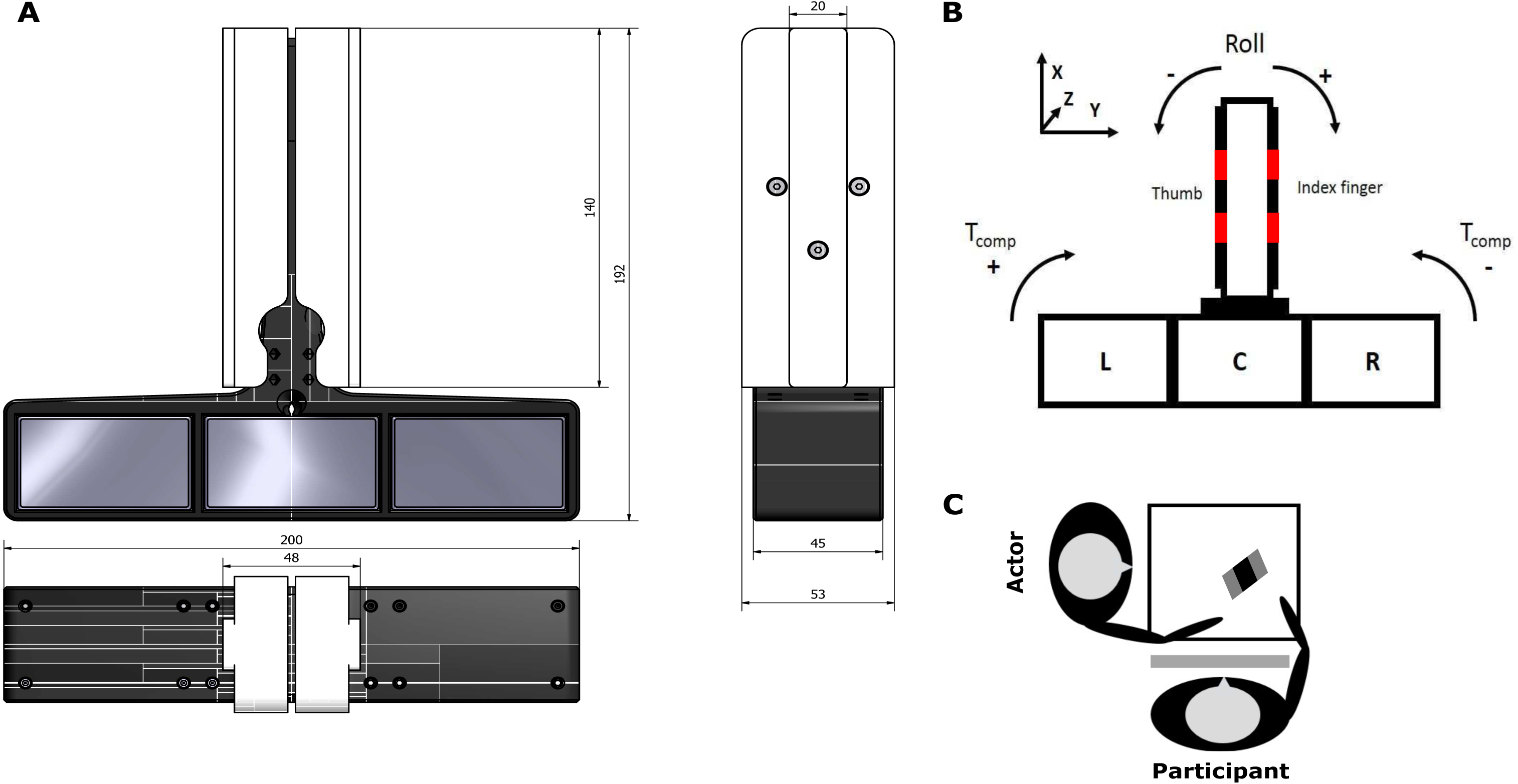
Experimental set-up. (**A**) Left: Frontal, side and top-down view of the ‘inverted T-shape’ manipulandum with dimensions (in mm). (**B**) Schematic drawing of the manipulandum with the three compartments indicated with L, C and R standing for left, central and right respectively. In red, the constrained locations are marked. X, Y and Z indicates the frame of reference for the force/torque sensors in the vertical component. (**C**) Dyadic positioning at a square table with the manipulandum placed in between the actor and participant. Figure modified from and reprinted with permission from Rens et al. (2020b)

### Data acquisition

For the present study, we used the same custom-built carbon fiber ‘inverted T-shape’ grip-lift manipulandum (Arsalis, Belgium; for all object dimensions see: Figure 1) as in one of our previous studies (Rens et al., 2021). The manipulandum consisted of a horizontal basis and a vertical block to which two 3D force/torque (F/T) sensors were attached. A cover plate (height × width: 140 × 53 mm) with a central protruding surface (height × width: 140 × 20 mm) was attached to each F/T sensor to block view on the sensors. On each protruding surface, we attached two pieces of fine-grained sandpaper (p600) (height x width: 20 x 20 mm). Both actor and participants were only allowed to place their fingertips on these constrained locations. The distance between the bottom side of the upper piece and the upper side of the lower piece of sandpaper on the same cover plate was 20 mm. As a result, the vertical distance between the center points of the same-sided pieces was 40 mm. This vertical distance was based on a preliminary investigation which showed that when lifting the asymmetrical weight distribution and keeping the basis horizontal, the load force difference between the two fingertips was zero if the fingertips were placed non-collinearly and distanced 40 mm from each other. The manipulandum’s horizontal basis was divided into three compartments enabling the placement of 3D-printed cuboids that were visually identical (height x width x depth: 55 x 35 x 40 mm). One cuboid was filled with lead particles and weighted 4.24 N, the other two were hollow and weighted 0.24 N each. Combined with the manipulandum, the total weight amounted to 8.67 N. The external torque (i.e., torque induced by the object’s weight distribution) could be changed by inserting the heavy cuboid in the left, center or right compartment and amounted to -245, 0 or + 245 Nmm, respectively (for the calculation of these values see Rens et al., 2021).

For collecting lifting-related parameters, we used two ATI mini-40 SI-40-2 F/T sensors (force range: 40, 40 and 120 N for x-, y- and z-axes respectively; force resolution: 0.01 N; torque range: 2 Nmm; torque resolution: 0.0005 Nmm) (ATI Industrial Automation, USA). F/T sensors were calibrated by the developer in accordance with the applicable QTI procedures. The maximum amount of error for the force and torque components were 1.50 % and 1.75 % respectively. Both F/T sensors were connected to a NI-USB 6221 OEM board (National Instruments, USA) which was connected to a personal computer. Data was acquired using a custom-written MATLAB script (Mathworks, USA) and sampled at 1 kHz.

### TMS procedure and EMG recording

#### General procedure

Electromyography (EMG) recordings were performed using Ag-AgCl electrodes which were placed in a typical belly-tendon montage over the right first dorsal interosseous muscle (FDI) and abductor pollicis brevis (APB). A ground electrode was placed over the processus styloideus ulnae. Electrodes were connected to a NL824 AC pre-amplifier (Digitimer, USA) and a NL820A isolation amplifier (Digitimer, USA) which in its turn was connected to a micro140-3 CED (Cambridge Electronic Design Limited, England). EMG recordings were amplified with a gain of 1000, high-pass filtered with a frequency of 3 Hz, sampled at 3000 Hz using Signal software (Cambridge Electronic Design Limited, England) and stored for offline analysis. For TMS stimulation, we used a (figure-of-eight; 70 mm) DuoMAG 70BF coil connected to a DuoMAG XT-100 system (DEYMED Diagnostic, Czech Republic).

For M1 stimulation, the coil was tangentially placed over the head to induce a posterior-anterior current flow and to elicit motor evoked potentials (MEPs) in both right FDI and APB. The optimal stimulation site (i.e., ‘hotspot’) was defined as the position from which MEPs with maximal amplitude were systematically recorded in both muscles. For finding the hotspot, we initially turned the stimulation intensity to 40% of the maximum stimulator intensity and increased the intensity step-wise while searching. The hotspot was marked on top of the scalp. Stimulation intensity (1 mV threshold) for each participant was defined as the lowest stimulation intensity that produced MEPs greater than 1 mV in both muscles and in at least four out of eight consecutive trials when stimulating at the predetermined hotspot (average stimulation intensity = 54 ± 5.6 % of maximum stimulator output). We assessed baseline (i.e., resting state) CSE before and after the experimental task. For this, participants received 12 TMS pulses at the previously defined stimulation intensity. During baseline assessment, participants were instructed to stay relaxed and keep their eyes open.

During the experiment, TMS was applied during both the actor (observation) and participant trials (execution). During actor trials, TMS was applied 300 ms after the actor lifted the object from the table (for definition of lift-off see: ‘Data analysis’). This timing was based on earlier work from our group (Rens et al., 2020) which showed clear motor resonance effects 300 ms after observed lift-off. In addition, similar studies have also applied TMS during the observed lifting phase to assess motor resonance effects during observation of object lifting (Alaerts, et al., 2010; Cretu et al., 2019; Senot et al., 2011). During participant trials, TMS was applied 400 ± 100 ms (jitter) after object presentation. As participants were instructed to only start reaching for the object after TMS was applied, the TMS stimulation was actually applied during lift planning. For participant trials (i.e., action execution) we decided to stimulate during planning, not execution, as we did not want to interfere with the participants’ lifting performance which would be caused by the unvoluntary muscle contraction due to M1 stimulation. TMS timing during participant planning was based on Loh et al. (2010). In their study, participants were instructed to lift a manipulandum with interchangeable. When a visual cue indicated the veridical object weight, CSE was differently modulated but only when TMS was applied 150 ms after object presentation. Considering that we used an object with a more complex property (i.e., interchangeable weight distribution), we decided to double the latency after which this TMS effect was present (i.e., 150 to 300 ms) to provide participants with enough time for planning. To ensure that participants could not anticipate the TMS timing, we decided to include a jitter of 100 ms We decided to use 300 ms as the lower threshold (thus 400 ms ± 100 ms jitter). To end, although TMS during lift planning was applied substantially later in our study than in the one of Loh et al. (2010), it should still elicit CSE modulation; Parikh et al. (2014) used a similar behavioral task and applied TMS at the ‘go cue’ which was given 1000 ms after the ‘task cue’. Briefly, in their study, CSE was also task-specifically modulated. As such, their findings indicate that TMS at the ‘go cue’ probes planning effects on CSE modulation, irrespective of the timing of the cue itself.

### Experimental set-up

#### Dyadic set-up

As shown in Figure 1C, participant and actor were comfortably seated at a square table with their lower arm resting on the table. The actor was seated on the left side of the participants so that the participant and actor were angled 90 degrees towards each other. The manipulandum was positioned between both individuals so both individuals could comfortably grasp and lift it. When grasping, both individuals were required to reach with their entire right upper limb causing their elbow to lift from the table. The manipulandum was distanced approximately 30 cm from each individual. In addition, participant and actor were asked to place their hand on a predetermined location in front of them to ensure consistent reaching throughout the experiment. It is important to note that the manipulandum was positioned as depicted in Figure 1 (i.e., targeted towards the participant). The manipulandum was rotated slightly (< 45 degrees) based on the participant’s preferences and to improve lifting comfort. Importantly, when the actor lifted the manipulandum it was positioned in this manner as well. When the actor reached for the manipulandum, their arm would move in front of the participant’s upper body (Figure 1C). Although this orientation likely occluded visual information about the left side of the manipulandum, we opted for this positioning rather than opposing both individuals for two reasons. First, Mojtahedi et al. (2017) showed that, when lifting an object together, individuals perform better when seated next to each other compared to opposed to each other. Second, Alaerts et al. (2009) showed that modulation of motor resonance is increased when observing actions from a first person point of view compared to a third person one. For our study, we argued that this side-by-side configuration would also enhance motor resonance effects in the participants. During the experiment, a switchable screen (MagicGlass) was placed in front of the participant’s face which was transparent during trials and returned opaque during inter-trial intervals. This screen blocked vision on the manipulandum when the experimenter would switch the cuboids between compartments, thus making participants blind to the weight distribution change. Last, one trial consisted of one lifting movement performed by either the actor or participant, thus being ‘participant trials’ or ‘actor trials’. Trial duration was 4 seconds and trial onset was indicated by the switchable screen turning transparent. Trial duration was based on preliminary testing and showed that individuals had sufficient time to reach, grasp, lift and return the object smoothly at a natural pace. Inter-trial interval was approximately 5 seconds during which the screen was opaque and the center of mass could be changed. To end, one (female) master’s student performed as the actor for all participants.

### Experimental procedure

#### General procedure

At the start of the session, participants gave written informed consent and were prepared for TMS (see ‘TMS procedure and EMG recording’). Afterwards, the experimenter explained the experimental task to the participants and gave the following instructions regarding the object lifting task: (1) lift the inverted T-shape to a height of approximately 5 cm at a smooth pace that is natural to you. (2) Only use thumb and index finger of the right hand and only place them on the sandpaper pieces. (3) You are required to use the same digit positioning the actor used in their preceding trial. (4) Keep the inverted T-shape’s base as horizontal as possible during lifting (i.e., ‘try to minimize object roll’). (5) The center of mass in your trials always matches the one in the actor’s preceding trial. In sum, the experimenter explained to the participants that they should try to minimize object roll during object lifting and that they could potentially rely on the observed lifting performance of the actor to plan their own lifts. Importantly, participants were instructed to always use the same digit positioning as the actor. This was done to ensure that we would have enough trials per experimental condition (see below).

After task instructions, participants were allowed to perform 3 practice lifts on the symmetrical weight distribution and 6 on each asymmetrical weight distribution (left or right). On half of the lifts with the asymmetrical weight distribution, participants were required to position their fingertips on the same height, i.e., ‘collinear positioning’. In the other half, they were required to place their fingertips ‘noncollinearly’, i.e., the fingertip on the heavy side was positioned higher than the fingertip on the light side (left asymmetrical: right thumb higher than right index; right asymmetrical: right thumb lower than right index). When lifting the symmetrical weight distribution, no compensatory torque should be generated. Accordingly, for this weight distribution participants were required to always place their fingertips collinearly as noncollinear positioning would cause participants to automatically generate compensatory torque. These practice trials were aimed to familiarize participants with the manipulandum.

#### Constrained digit positioning

Fu et al. (2010) showed that appropriate compensatory torque can be generated by many valid digit position-force coordination patterns. As motor resonance is driven by movement features such as fingertip forces, we wanted to limit the digit positioning options in order to reduce potential variability in motor resonance effects. We argued that constrained digit positioning and limited experimental conditions would enable us to find more reproducible results. As mentioned before, noncollinear positioning was defined based on the vertical difference between fingertips which caused the load force difference to approximate zero. Collinear positioning was included as we considered it a ‘neutral’ condition to contrast the noncollinear one. Because the fingertip on the heavy side has to generate more force when positioned collinearly compared to noncollinearly, we argued that these two conditions would allow us to disentangle whether the observer’s motor system would encode observed positioning or observed forces. When using collinear digit positioning, irrespective of the weight distribution, actor and participant were required to place their fingertips on the upper sandpaper pieces on each side (Figure 1B). When using noncollinear digit positioning for the asymmetrical weight distribution, the finger on the heavy side was placed on the upper piece whereas the finger on the light side was placed on the lower piece. For instance, when the heavy cuboid was inserted in the left compartment, the thumb and index finger were placed on the upper and lower piece of their respective side (Figure 1). The actor was instructed to use these digit positioning strategies which ensured that participants would use the same digit positionings. We opted for these digit positionings (primarily collinear positioning on the upper pieces instead of the lower ones) to protect the EMG electrodes over the thumb muscle from rubbing over the manipulandum’s base.

#### Experimental task

After task instructions, participants performed the object lifting task with the actor. The actor would change the inverted T-shape’s center of mass and verbally declare that she would execute the next trial. Verbal declaration took place before switching of the cuboids. After trial completion by the actor, participant performed 3 back-to-back trials with the same weight distribution and using the same digit positioning as the actor. We decided to have participants perform 3 repetitions based on Fu et al. (2010). To ensure that participants could not rely on sound cues potentially indicating the new weight distribution, the experimenter always removed and replaced all 3 cubes after randomly rotating the inverted T-shape prior to each actor trial. These actions were never done before participant trials as they were explained that the center of mass in their trials would always match the one of the actor’s preceding trial.

During the object lifting task, the experimenter and participants performed 20 transitions from the middle compartment (i.e., symmetrical weight distribution) to each side (i.e., asymmetrical weight distributions). When the experimenter lifted the new asymmetrical weight distribution (e.g., left-sided), she would use collinear positioning in 10 lifts and noncolinear positioning in the other 10. Trial amount per experimental condition was based on Senot et al. (2011) who used 10 trials per TMS condition when investigating motor resonance effects during lift observation. After 4 trials were performed on the asymmetrical weight distribution (i.e., 1 actor and 3 participant trials), the experimenter would change the object’s weight distribution again. Most of these transitions were ‘normal’ and consisted of the actor changing the heavy cuboid back to the middle compartment for washing out the internal representation for the asymmetrical weight distribution. However, 10 transitions were ‘catch transitions’ in which the asymmetrical weight distribution was first changed to the other side (e.g., left to right asymmetrical) and only then to the middle compartment to wash out the internal representation for asymmetrical. We included these catch transitions to ensure that participants would not anticipate the typical change from asymmetrical to symmetrical even though we still wanted to wash out the internal representation for asymmetrical. Catch transitions were inserted equally after each lifting sequence on each asymmetrical weight distribution. Considering that we had a 2 (side: left or right asymmetrical) by 2 (positioning: collinear or noncollinear) design when lifting the asymmetrical weight distribution, either 2 or 3 catch transitions were performed after each lifting sequence (e.g., 2 catch transitions after the lifting sequence with collinear positioning on the right asymmetrical weight distribution). The actor randomly decided which digit positioning to use on the new asymmetrical weight distribution in the catch transitions. Last, in both the normal and catch transition to symmetrical, the standard amount of trials (1 actor and 3 participant’s trials) were performed after each weight distribution change.

The object lifting task was split over 4 experimental blocks with a short break between blocks. For each participant, transition order was pseudo-randomized within each experimental block. As such, each experimental block contained 5 transitions to left and right asymmetrical. In addition, either 2 or 3 of these transitions to each side were performed with one digit positioning type (e.g., collinear) and the other 3 or 2 transitions with the other digit positioning type (e.g., noncollinear). Transitions to a specific side (e.g., symmetrical to left-sided asymmetrical) would repeat maximally 2 times back-to-back and catch transitions were spread equally over all blocks. The full experimental session lasted approximately 2 h.

### Data analysis

#### Behavioral data

Data collected with the F/T sensors were sampled in 3 dimensions at 1000 Hz and smoothed using a fifth-order Butterworth low-pass filter (cut-off frequency: 15 Hz). For each sensor, grip force (GF) and load force (LF) were defined as the exerted force perpendicular to the normal force (Y-direction on Figure 1) and the exerted force parallel to the normal force (X-direction on Figure 1), respectively. Digit positioning was defined as the vertical coordinate (X-direction on Figure 1) of the fingertip’s center of pressure on each cover plate. The center of pressure was calculated from the force and torque components measured from the respective F/T sensor relative to its frame of reference, using formula 1.

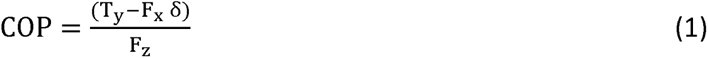

In formula 1, COP = center of pressure, T_y_ = Torque in the Y-direction, F_x_ = Force in the X-direction, F_z_ = Force in the Z-direction, δ = cover plate thickness (1.55 mm). Compensatory torque was defined as the net torque generated by an individual to offset the external torque caused by the object’s weight distribution and was calculated with formula 2 (we refer the reader to the supplementary materials of Fu et al., 2010 for the detailed explanation of the formula).

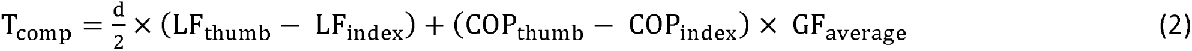

In formula 2, T_comp_ = Compensatory torque, d = horizontal distance between the digits (48 mm; Figure 1; Y-direction), LF_thumb/index_ = Load force generate by the thumb and index finger, respectively, COP_thumb/index_ = center of pressure of the thumb and index finger, respectively, GF_average_ = averaged amount of GF exerted by the thumb and index finger.

To investigate the effects of lift observation on the performance of the participants we used the following variables: Digit positioning difference, defined as the difference between the COP of the thumb and the index finger (positive values indicate a thumb placement higher than that of the index finger), compensatory torque, total grip force, and load force difference, defined as the difference between load forces generated by the thumb and index finger (positive values indicate the thumb generating more load force than the index finger). We included difference in digit positioning to investigate whether actor and participants placed their fingertips appropriately. However, it is important to note that during data processing we excluded trials (both force and TMS data) in which fingertips were not placed on the sandpaper pieces (< 1% of all trials). We considered compensatory torque as our key indicator of performance as it results from the combination of grip and load forces as well as digit positioning and because we explicitly asked participants to minimize object roll during lifting (‘task goal’). Moreover, Fu et al. (2010) showed a strong linear correlation between compensatory torque and peak object roll, arguing its validity as our key indicator of performance. We included total grip force and load force difference to explore their potential effect on CSE modulation during observation and planning. Last, it is important to note that, for analysis purposes, we inversed the sign for compensatory torque for the right asymmetrical weight distribution, which allows for better comparisons regarding performance between the left and right side. We argued not to do so for load force and digit position differences for interpretability with respect to motor resonance effects.

In line with Fu et al. (2010), we extracted digit positioning difference at early object contact, which we defined as total GF > 1 N. Considering that we had no proxy of lift onset (again see Fu et al. 2010), we decided to approximate lift onset by using object lift-off instead. In line with our previous work (Rens et al., 2020, 2021; Rens & Davare, 2019), we defined lift-off as the time point where total load force > 0.98 x object weight. As such, we extracted total grip force, load force difference and compensatory torque at object lift-off.

*EMG data*. From the EMG recordings, we extracted the peak-to-peak amplitude of the MEP using a custom-written MATLAB script. All EMG recordings were visually inspected and afterwards analyzed in the script. Trials were excluded when the MEP was visibly contaminated by noise (i.e., spikes in background EMG) or when an automatic analysis found that background EMG was larger than 50 µV (root-mean-square error) in a time window of 200 ms prior to the TMS stimulation. In line with previous work of our group (Rens et al., 2020) we excluded outliers for each participant separately. Outliers were defined as values exceeding the mean ± 3 SD’s. The total amount of removed MEPs was 1.68 %.

For each participant, all MEPs collected during the experimental task were z-score normalized for observation and planning separately. Last, we also assessed pre-stimulation (‘background’) EMG by calculating the root-mean-square across a 100ms interval ending 50ms prior to TMS stimulation. To end, we did not normalize baseline CSE measurements to allow for direct comparisons between muscles.

### Statistical analyses

For statistical purposes, we did not include catch trials (center of mass change from side to side) and trials in which the weight distribution changed from asymmetrical to symmetrical. First, we excluded transitions to the symmetrical weight distribution as we had less than 10 MEPs for these conditions. As mentioned before, we had 20 transitions to each asymmetrical weight distribution, 10 in which the actor used collinear positioning and 10 in which she used noncollinear positioning. After performing this lifting sequence, the weight distribution normally changed to symmetrical. However, as mentioned before, we also included 10 catch transitions in which one asymmetrical weight distribution changed to the other (i.e., 5 changes from left to right and 5 from right to left). Due to these 10 catch transitions, we had less than 10 MEPs for each transition to symmetrical. For instance, when a lifting sequence was performed on the right asymmetrical weight distribution with noncollinear digit positioning only 7 or 8 normal transitions were performed in which the weight distribution changed directly back to symmetrical. Given the limited amount of MEP data for these changes with respect to relevant motor resonance studies (Alaerts et al. 2010; Buckingham et al. 2014; Senot et al. 2011) and the actor choosing her digit positioning randomly in these catch transitions, we decided to not include these trials for analysis purposes. Note that we included lifts for the symmetrical weight distribution lifts in the analysis of the actor’s performance, to compare this against the asymmetrical distributions. Second, we excluded catch trials for analysis purposes due to the very limited amount we had.

We investigated whether the actor’s and participants’ performance changed between experimental blocks. Considering we did not find any learning effects, we collapsed our data across experimental blocks. All statistical analyses were performed in SPSS statistics version 25 (IBM, USA) and are described below. For each parameter of interest, we performed a separate analysis. We used linear mixed models (LMM) considering our coding for the noncollinear condition (left noncollinear: thumb higher than index; right noncollinear: thumb lower than index). This allowed to consider digit positioning to be a ‘nested’ factor within the weight distribution for our analyses (see below) instead of a ‘main’ factor as in a typical ANOVA. The reason hereof is that it has been shown that body representations have different preferential associations between the fingers and their positioning in space (e.g., index finger on top and thumb on the bottom) (Romano et al., 2017). As such, it is plausible that these preferential associations modulate CSE, recorded in the FDI (index finger) and APB (thumb) muscles, differently during lift planning and observation.

For the actor’s data (behavioral only), we used the factors SIDE (mid, left and right) and POSITION (collinear and noncollinear which were coded as described in the preceding paragraph). We included the actor’s collinear lifts on the symmetrical weight distribution (i.e., mid) to investigate changes in lift performance. Specifically, if the actor correctly planned to lift an asymmetrical weight distribution, her lift performance should differ significantly from that when she planned to lift the symmetrical weight distribution (Rens and Davare 2019). We included SIDE as a main effect and nested POSITION within SIDE (i.e., POSITION_SIDE_).

For the participants’ data (behavioral and MEPs), we used the factors SIDE (left and right only), POSITION (collinear and noncollinear) and REPETITION (first, second and third lift after the center of mass change). Please note that the factor REPETITION and all its interaction effects described below were only included during lift planning, not observation, which we investigated separately. SIDE and REPETITION were included as main effects. Again, POSITION was nested within SIDE. For the participants’ data we also included the interaction effect SIDE X REPETITION as well as REPETITION X POSITION_SIDE_. Here, we did not include the lifts on the symmetrical weight distribution as lift performance of participants can be investigated by comparing lifts without tactile feedback (i.e., first lift after lift observation) to their own lifts on the same object with tactile feedback (second and third lifts). As such, lift performance is quantified by comparing lifts without and with tactile feedback (Fu et al. 2010; Reichelt et al. 2013; Rens and Davare 2019). To analyze motor resonance effects during lift observation, we used the factors SIDE (left and right) and POSITION (collinear and noncollinear). We included SIDE as a main effect and nested POSITION within SIDE (i.e., POSITION_SIDE_).

For both the actor and participant data, we included the intercept in the model. Moreover, for all participants’ analyses we also included the participant’s number as random effect. Last, we decided to include the mixed model covariance structures as first-order autoregressive based on the assumption that correlation in residuals between factor levels was identical across levels. We used type III sum of squares and Maximum Likelihood (ML) for mixed model estimation and Bonferroni corrections for pairwise comparisons. All data in text is presented as the mean ± SEM. P-values < 0.05 are discussed as statistically different.

**Figure 2.**
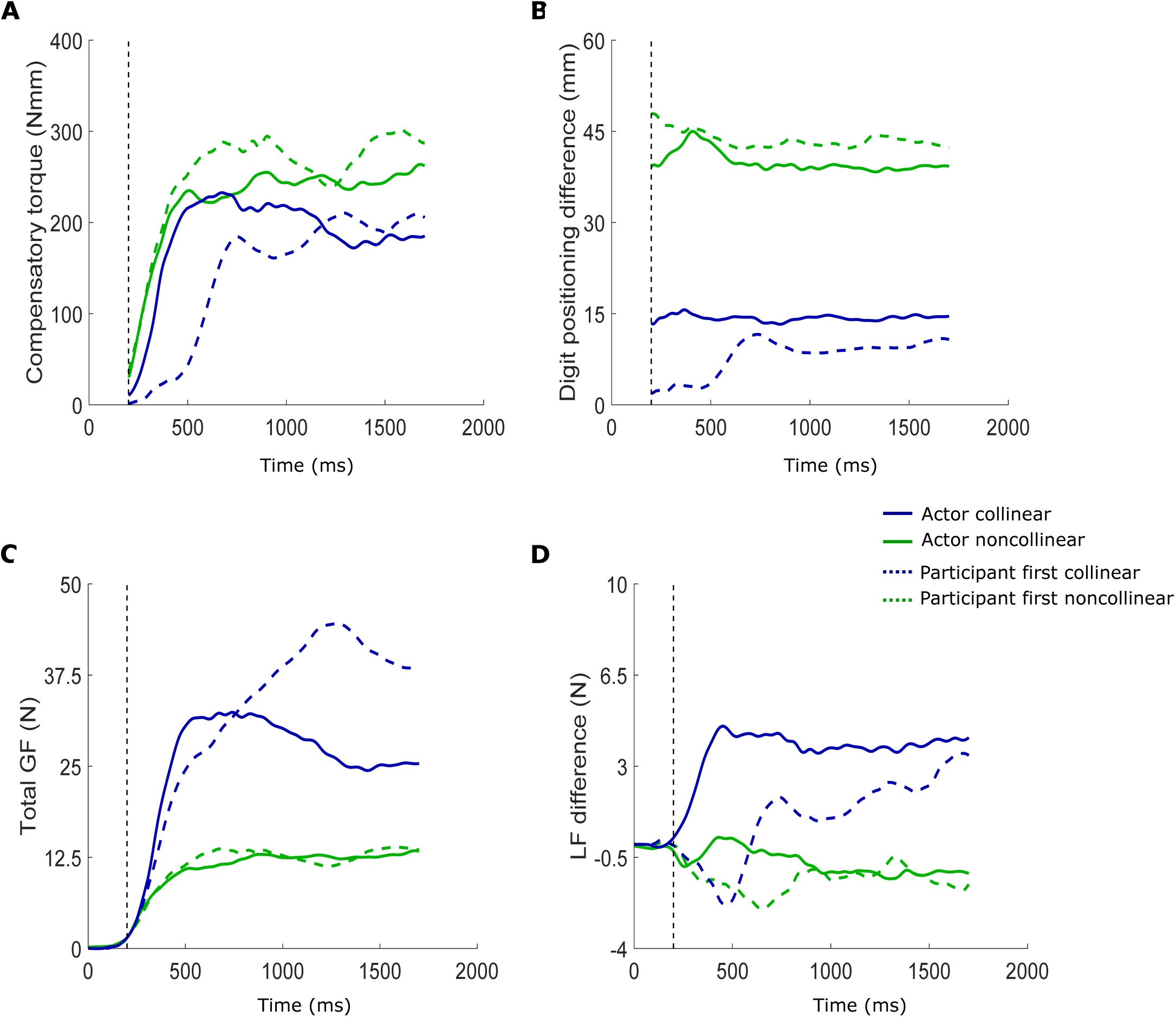
E**xample parameter traces.** One lift example of performance on a left-sided asymmetrical weight distribution for the actor (solid traces) and first lift of a participant (dashed traces). These traces show the typical evolution of different parameter profiles over time for a lift with fingertips positioned on the same height (‘collinear’; in blue) or with fingertips positioned on different heights (‘noncollinear’; in green). (**A**) Compensatory torque (in Nmm). (**B**) Digit positioning difference, i.e., vertical height difference in center of pressure of the fingertips (position thumb - position index finger; in mm). (**C**) Total grip force (in N). (**D**) Load force difference (load force thumb – load force index finger; in N). Vertical dashed line on each figure indicates object contact. We cleared compensatory torque and digit positioning difference values before object contact as they are highly contaminated by noise.

## Results

In the present study, we investigated how CSE is modulated when observing and planning lifts of objects with an asymmetrical weight distribution. Participants performed an object lifting task in turns with an actor and were required to lift a manipulandum with interchangeable center of mass as skillfully as possible, i.e., minimize object roll by generating appropriate compensatory torque. When participants were required to lift an object with unknown weight distribution, they first observed the actor lift the new weight distribution. Afterwards, the participants lifted this weight distribution three times. Importantly, participants were instructed to place their fingertips on the same constrained locations the actor used. After participants performed their three lifts, the actor changed the weight distribution and again lifted the new weight distribution. Accordingly, participants could potentially derive critical information about the object’s weight distribution by observing the actor’s lifts. During the behavioral task, TMS was applied during observed lifting (after observed lift-off) and during lift planning in the actor and participant trials, respectively.

Traces of the included lifting parameters, when lifting an asymmetrical weight distribution, can be found in Figure 2, showing the difference between a single lift of the actor (solid lines) with collinear (blue) and noncollinear positioning (green). In addition, the dashed traces represent a single first lift of a participant after observing the actor’s lift. As the actor could anticipate the object’s weight distribution, she would predictively generate compensatory torque to minimize object roll (Figure 2A; blue and green solid traces). When the participants lifted the object after observing the actor, they were able to generate appropriate compensatory torque after observing noncollinear positioning (green dashed trace) but not after observing collinear one (blue dashed trace). Accordingly, the difference in lift performance can be quantified through differences in compensatory torque. As mentioned before, appropriate compensatory torque is the resultant of a valid digit positioning – force coordination pattern (Fu et al., 2010). For instance, it can be seen in Figure 2B that in the actor’s (blue solid) and participant’s lift (blue dashed), fingertips were positioned similarly. However, as the actor but not the participant generated appropriate compensatory torque, it can be seen in Figure 2D that the load force difference increased faster for the actor than the participant. Conversely, when lifting with noncollinear digit positioning (green traces), the fingertip on the heavy side is positioned higher than the fingertip on the light side (Figure 2B). Due to this vertical height difference, the fingertip on the heavy side does not have to generate more load force than the fingertip on the light side (Figure 2D), resulting in a load force difference close to zero. Last, as can be seen in Figure 2C, the fingertips generate more grip force when positioned collinearly compared to noncollinear positioning.

**Figure 3.**
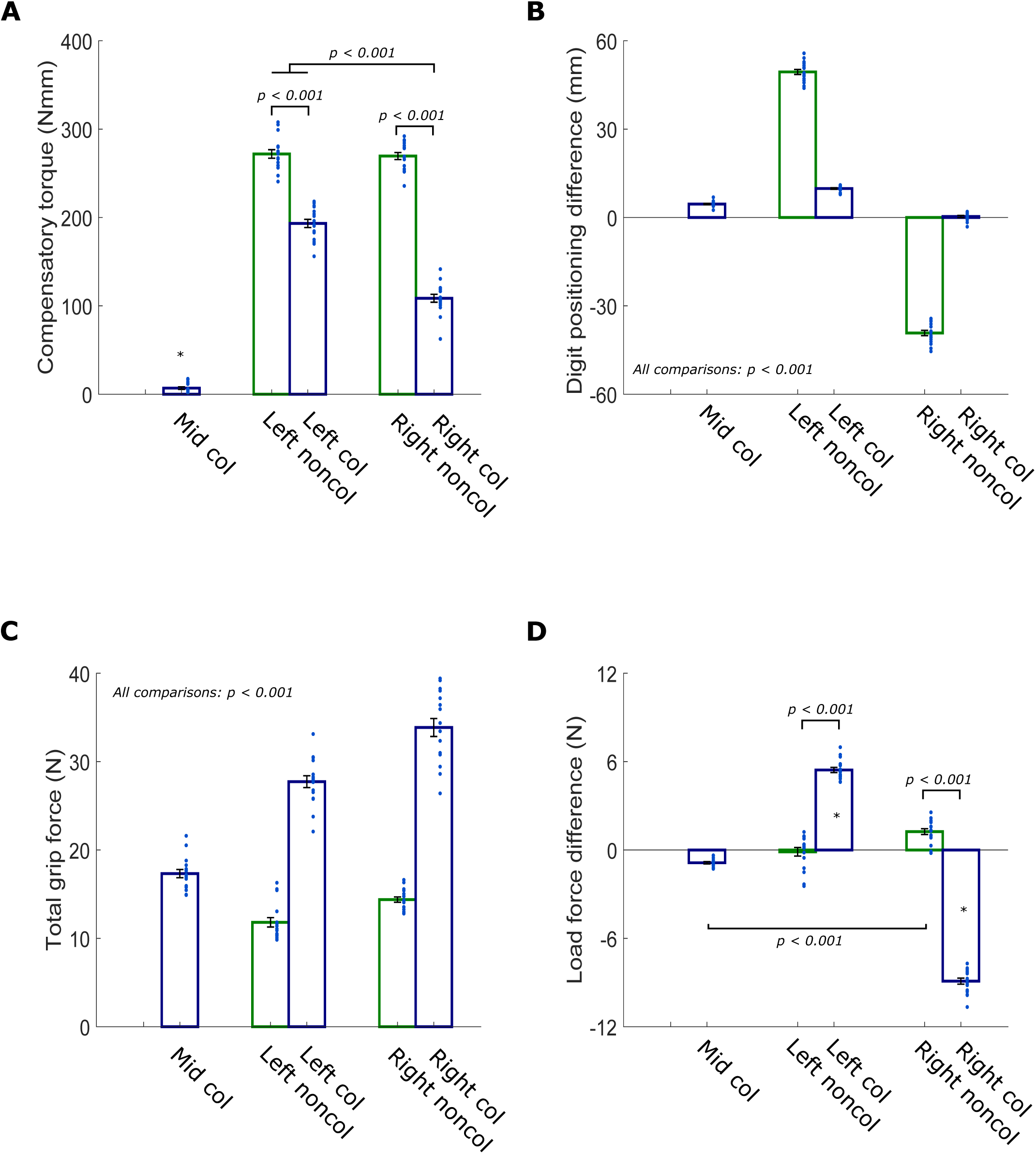
Lift performance of the actor. The actor’s averaged lift performance. The actor used either collinear (col; blue) or noncollinear (noncol; green) digit positioning to the object. Each scatter dot represents the averaged performance of the actor as observed by one participant. (**A**) Compensatory torque (in Nm). (**B**) Digit positioning difference (in mm). (**C**) Total amount of grip force (in N). (**D**) Load force difference (in N). All data is presented as the mean ± SEM.

### Actor’s lift performance

The actor’s lift performance should have been consistent due to (a) our constrained positioning conditions and (b) her changing the weight distribution. However, we decided to analyze her performance for verification and to quantify which observed movement features might have driven motor resonance effects. We expected that the actor would comply with task instructions and would always lift the manipulandum skillfully (i.e., generate appropriate compensatory torque).

#### Compensatory torque at lift-off

As the actor was not blinded to the weight distribution change, she generated significantly more compensatory torque when lifting the asymmetrical weight distributions (left = 232.56 ± 4.22 Nmm; right = 189.03 ± 3.39 Nmm) compared to the symmetrical one (mean = 6.90 ± 1.38 Nmm; both *p* < 0.001) (SIDE: F_(2,80)_ = 1135.70; *p* < 0.001). Although the actor was instructed to lift the asymmetrical weight distributions as skillfully as possible, she failed to do so when using collinear positioning: as shown in Figure 3A, the actor generated more compensatory torque when lifting asymmetrical weight distributions with noncollinear positioning (between adjacent bars: each *p* < 0.001) (POSITION_SIDE_: *F*_(2,80)_ = 518.184; *p* < 0.001). Noteworthy, the performance when lifting noncollinearly was similar between asymmetrical weight distributions (Figure 3A; between green bars: p = 1.00). Last, the actor’s lift performance was the least skillful when collinearly lifting the right asymmetrical weight compared to all other asymmetrical conditions (*all p* < 0.001).

#### Digit positioning at early object contact

Although digit positioning was constrained to small sandpaper pieces and we removed trials in which fingertips were not positioned correctly, we provide these results for clarity. Please note that positive values for difference in digit positioning indicate that the thumb was positioned higher than the index finger and negative values indicate that the index finger was positioned higher than the thumb. Briefly, when the actor was instructed to use noncollinear positioning, she appropriately placed her fingertips further apart compared to the collinear conditions (Figure 3B; green compared to blue bars: all *p* < 0.001) and differently than the other noncollinear condition (between green bars: *p* < 0.001) (POSITION_SIDE_: *F*_(2,80)_ = 2342.74, *p* < 0.001). Noteworthy, digit positioning also differed significantly between all collinear conditions (*all p* < 0.001) which indicates that the weight distribution caused subtle differences in the collinear digit positioning. However, considering how small the differences between collinear conditions are (Figure 3B; Mid col = 4.55 ± 0.24 mm; Left col = 9.86 ± 0.25 mm; Right col = 0.32 ± 0.36 mm), it is unknown to which extent they were visible to the participants.

#### Total grip force at lift-off

When the actor lifted the asymmetrical weight distributions noncollinearly, she used less grip force than when lifting all weight distributions collinearly (Figure 3C; green compared to blue bars: all *p* < 0.001) (POSITION_SIDE_: *F*_(2,80)_ = 405.68; *p* < 0.001). Importantly, differences between all conditions were significant (*all p* < 0.001) suggesting that the actor used a unique amount of grip force in each condition. It also important to note that the actor used, on average, more grip force when lifting the right asymmetrical weight distribution (mean = 24.12 ± 0.63 N) compared to the left one (mean = 19.77 ± 0.52 N; *p* < 0.001) (SIDE: F_(2,80)_ = 46.13; *p* < 0.001).

**Figure 4.**
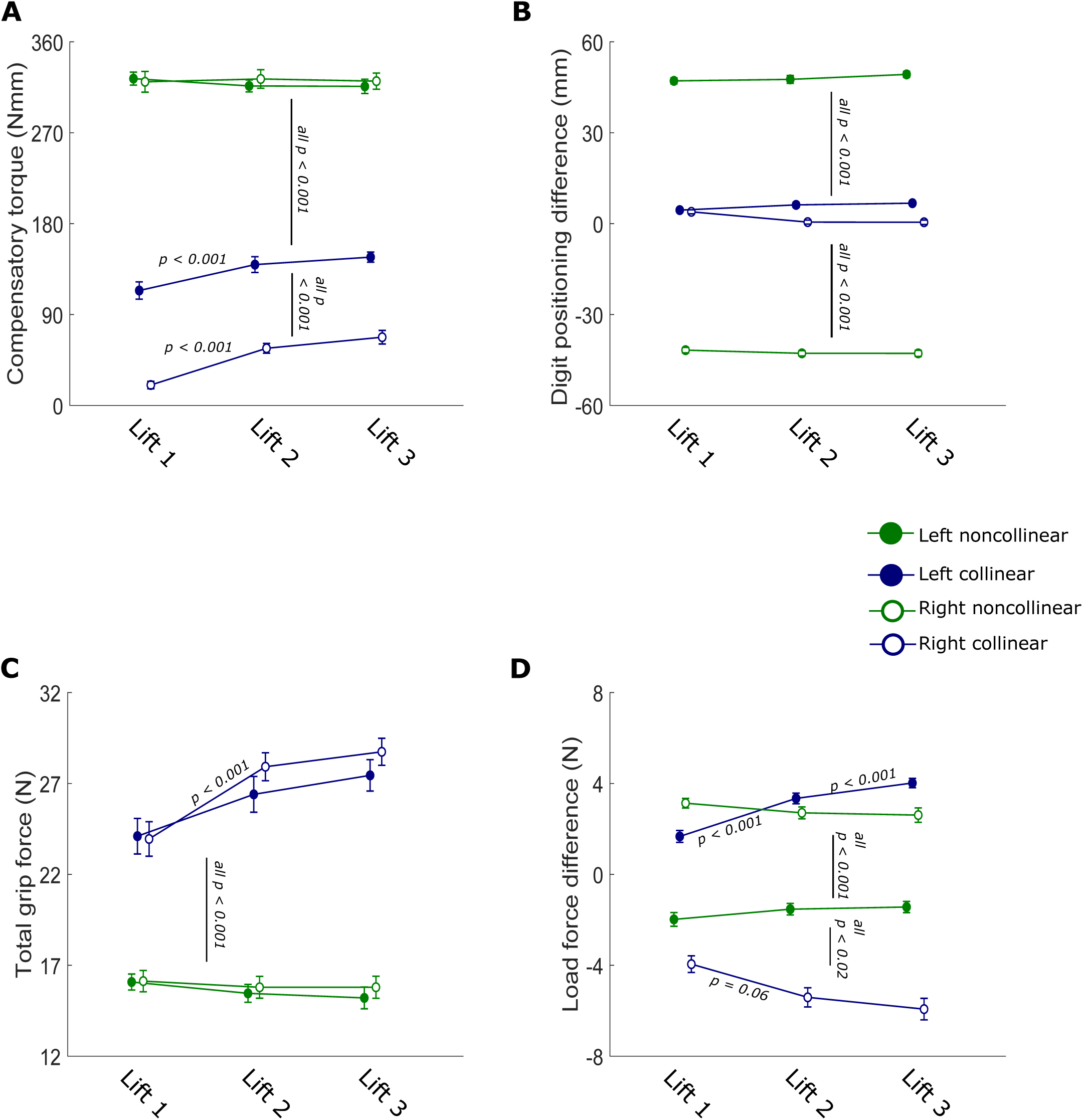
Lift performance of the participants. Connected scatterplot showing the averaged lift performance of the participants for their three lift repetitions. Performance is shown for when the weight distribution was left (solid scatter) or right (empty scatter) asymmetrical. Participants used either collinear (blue) or noncollinear (green) digit positioning for lifting. (**A**) Compensatory torque (in Nmm). (**B**) Digit positioning difference (in mm). (**C**) Total grip force (in N). (**D**) Load force difference (in N). All data is presented as the mean ± SEM. Vertical lines with a p-value indicate that all scatters above this line differ significantly from all scatters below. P-values between two connected scatters indicate that those two differ significantly from each other. For within-condition comparisons, only significant differences between two subsequent trials are shown (e.g., between trial one and two but not between trial one and three).

#### Load force difference at lift-off

Please note that these values are expressed as the difference in load force between the thumb and index finger therefore positive values indicate that the thumb exerted more load force than the index finger and negative values indicate that the index finger exerted more load force than the thumb. First, when the actor lifted the asymmetrical weight distributions collinearly, she generated significantly more load force with the fingertip on the heavy side (left = 5.43 ± 0.18 N; right = -8.91 ± 0.20 N) compared to lifting the same weight distribution noncollinearly (Figure 3D; between adjacent bars: *p* < 0.001) and compared to all other conditions (*all p* < 0.001) (POSITION_SIDE_: *F*_(2,80)_ = 877.52; *p* < 0.001). Second, we found no evidence that the load force difference was different when the actor lifted the asymmetrical weight distributions noncollinearly (Figure 3D; between green bars: p = 0.09).

### Participants’ lift performance

We included the participants’ lifting parameters to investigate their lift performance after lift observation (first lift) and haptic feedback (second and third lifts) and how it might have affected CSE modulation during lift planning. Based on our previous work (Rens et al., 2021), we expected that lift performance would be suboptimal after lift observation. That is, participants would perform better in their second and third lifts compared to their first ones.

#### Compensatory torque at lift-off

Our results revealed that there were large differences in lift performance between digit positioning conditions. When participants used noncollinear positioning, they overcompensated for the asymmetrical weight distribution and generated more compensatory torque (left = 319.50 ± 5.68 Nmm; right = 321.57 ± Nmm; Figure 4A green lines) than required (245 Nmm). In contrast, when lifting collinearly, participants could not generate appropriate compensatory torque (left = 133.35 ± 6.73 Nmm; right = 48.14 ± 4.14 Nmm; Figure 4A blue lines). Although lift performance did not differ between the noncollinear conditions (p = 1.00), it did differ between noncollinear and collinear ones (*all p* < 0.001). In addition, participants performed worse when collinearly lifting the right asymmetrical weight distribution compared to all other conditions (all < 0.001) (POSITION_SIDE_: *F*_(2,179)_ = 1571.60; *p* < 0.001). When participants lifted the asymmetrical weight distributions noncollinearly, they did not generate more compensatory torque over repetitions (between green connected scatters: all p = 1.00). This indicates that their lift performance did not differ significantly after lift observation (first repetition: left = 323.56 ± 6.44 Nmm, right = 320.41 ± 10.26 Nmm) and after having tactile feedback (second repetition: left = 316.34 ± 5.86 Nmm; right = 323.23 ± 9.22 Nmm; third repetition: left = 315.89 ± 7.07 Nmm; right = 321.07 ± 7.99 Nmm) (REPETITION X POSITION_SIDE_: *F*_(4,179)_ = 4.99; *p* < 0.001). In contrast, when using collinear positioning, participants performance improved over repeated lifts. This increase was significant from their first (left = 113.78 ± 8.57 Nmm, right = 20.26 ± 3.71 Nmm) to second lift (left = 139.44 ± 7.84 Nmm, right = 56.60 ± 4.82 Nmm) (*both p* < 0.009), but not significant from their second to third lift (left = 146.83 ± 5.02 Nmm, right = 67.57 ± 6.69 Nmm) (both p > 0.54) (Figure 4A blue connected scatters). To end, when participants used collinear positioning, the difference in compensatory torque was also statistically different between the first and third lifts for each side (left: p = 0.005; right: *p* < 0.001).

#### Digit positioning at early object contact

As participants were instructed to imitate the actor’s constrained digit positioning and we excluded trials with incorrect digit positioning, we primarily report digit positioning for transparency reasons. As shown in Figure 4B, participants did not change their digit positioning over the multiple repetitions within each condition (REPETITON X POSITION_SIDE_: *F*_(4179)_ = 0.75; p = 0.56). This indicates that they adhered to the constrained digit positionings. Logically, when lifting the left asymmetrical weight distribution noncollinearly, participants placed their thumb higher than their index finger (mean = 48.22 ± 0.1.06 mm) and vice versa for the right asymmetrical weight distribution (mean = -42.45 ± 0.84 mm; *p* < 0.001). These noncollinear positionings also differed significantly from those when lifting collinearly (*all p* < 0.001) (POSITION_SIDE_: *F*_(2,179)_ = 3605.93; p< 0.001). Noteworthy, collinear digit positioning for the left (mean = 5.80 ± 0.67 mm) and right (mean = 1.62 ± 0.54 mm) asymmetrical weight distributions also differed significantly (p = 0.003; effect of POSITION_SIDE_). Although this difference is significantly different, it is unlikely that participants planned their digit positioning differently between sides in the collinear condition as this difference is only ± 4 mm and the constrained locations for digit positioning were only 20 mm wide. Arguably, it is more likely that this small difference between sides is driven by skin deformation due to external torque caused by the weight distributions (Kalra et al., 2016) or by slipping of the fingertips (Johansson & Westling, 1984).

#### Total grip force at lift-off

As shown by the green line plots in 4C, participants scaled their grip forces similarly when using noncollinear positioning for both weight distributions and all repetitions (all p = 1.00) (REPETITION X POSITION_SIDE_: *F*_(4,179)_ = 5.53; *p* < 0.001). In contrast, this lifting consistency was lower when using collinear digit positioning. When participants lifted the right asymmetrical weight distribution collinearly, participants significantly increased their grip forces from their first lift (mean = 23.95 ± 0.95 N) to their second one (mean = 27.92 ± 0.77 N; *p* < 0.001) but not from their second to third one (mean = 28.74 ± 0.74 N; p = 1.00). The difference between the first and third lift was also statistically different (p < 0.001). When lifting the left asymmetrical weight distribution collinearly, participants did not significantly increase their grip force from their first (mean = 24.10 ±0.98 N) to second lift (mean = 26.40 ± 0.98 N; p = 0.51) and also not from their second to third lift (mean = 27.45 ± 0.86 N; p = 0.86). However, the amount of grip force participants generated in this condition was statistically different between the first and third lift (p = 0.05). By and large, these findings indicate that when participants lifted the asymmetrical weight distributions collinearly, they increased their grip force over the three repetitions. To end, as shown in Figure 4C, the amount of grip forces applied when lifting collinearly or noncollinearly differed significantly for both weight distributions and all repetitions (*all p* < 0.001).

#### Load force difference at lift-off

Based on our experimental set-up and constrained positioning, we expected that noncollinear positioning would lead to a load force difference close to zero. Conversely, collinear positioning would lead to the fingertip on the heavy side scale its load force larger than the finger on the light side. As shown in Figure 4D, when participants used noncollinear positioning, the load force difference was significantly different for the left (mean = -1.65 ± 0.24 N) and right (mean = 2.82 ± 0.24 N; *p* < 0.001) asymmetrical weight distributions (POSITION_SIDE_: *F*_(2,179)_ = 667.79; *p* < 0.001). This difference was not only present during their first lift (left = -1.98 ± 0.30 N; right = 3.13 ± 0.21 N) but persisted during their second (left = -1.53 ± 0.25 N; right = 2.71 ± 0.26 N) and third lifts (left = -1.44 ± 0.25 N; right = 2.61 ± 0.32 N) (between sides: all *p* < 0.001) (REPETITION X POSITION_SIDE_: *F*_(4,179)_ = 3.69; p = 0.006). Furthermore, our results provide no evidence that participants, in the noncollinear condition, scaled their load forces differently over multiple repetitions for each side (between repetitions for each side: all p = 1.00). As such, our results do not indicate that participants altered their load force difference based on tactile feedback.

In line with our expectations, our results show that when participants used collinear positioning, they exerted more load force with the finger on the heavy side (left = 3.01 ± 0.21 N; right = -5.10 ± 0.38 N; between sides: *p* < 0.001). Moreover, when using collinear positioning for each weight distribution, the load force difference became larger from the first (left = 1.67 ± 0.26 N; right = -3.95 ± 0.37 N) to the second lift (left = 3.34 ± 0.23 N; right = -5.41 ± 0.42 N). Noteworthy, this difference was significant for the left asymmetrical weight distribution (p < 0.001), but not for the right asymmetrical one (p = 0.06). These findings indicate that participants altered their load force difference based on tactile feedback, in particular, when lifting the left asymmetrical weight distribution. This effect further increased from the second to third lift for the left (mean = 4.02 ± 0.21 N, *p* < 0.001) but not right (mean = -5.93 ± 0.47 N, p = 0.26) asymmetrical weight distribution. To end, the difference between the first and third lift for each side was also statistically different (left: *p* < 0.001; right: p = 0.01).

**Figure 5.**
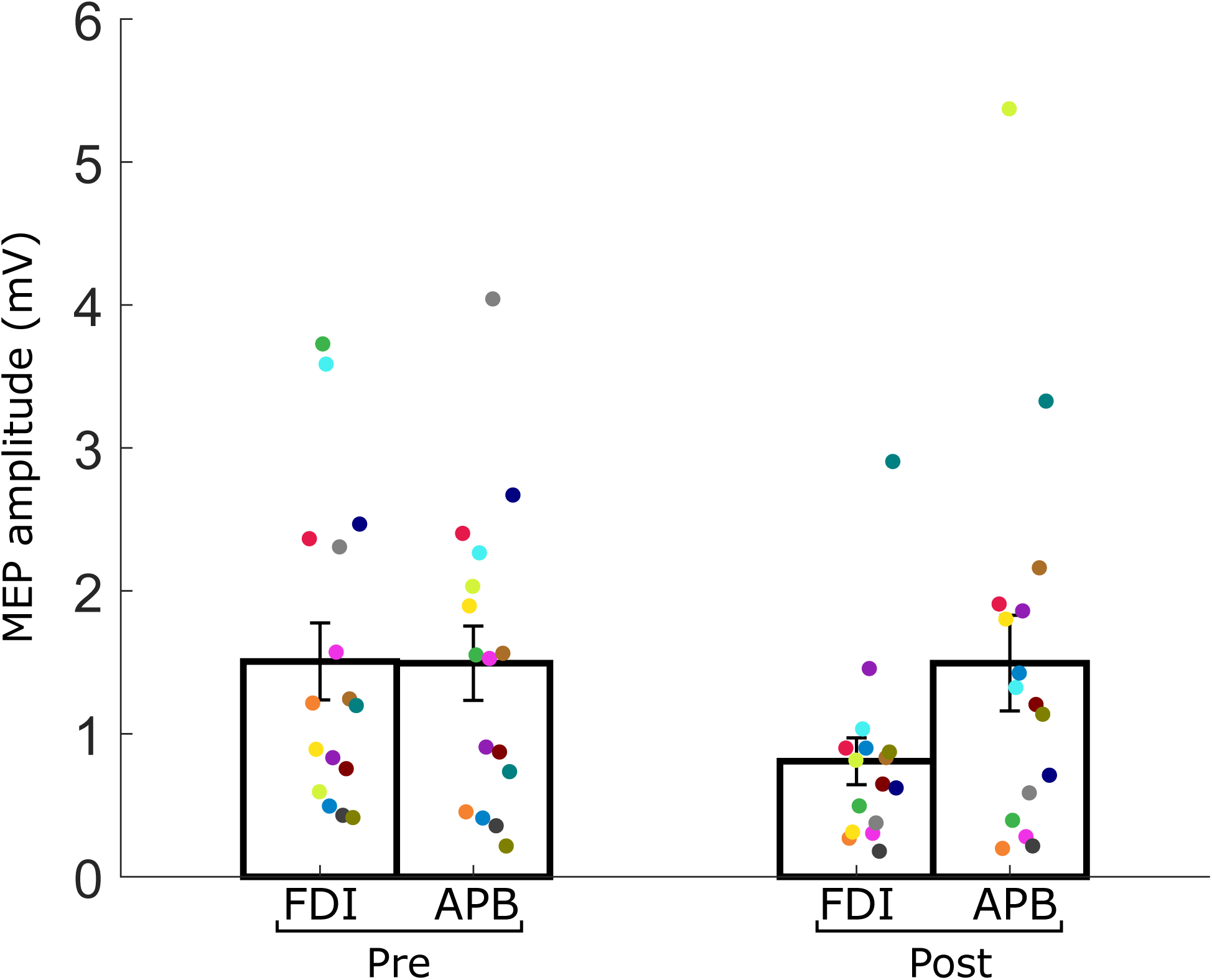
Corticospinal excitability during baseline (resting state). Average motor evoked potential (MEP) values in mV during baseline before (pre) and after (post) the experiment. MEPs were recorded from first dorsal interosseous muscle (**FDI**) and from the abductor pollicis brevis muscle (**APB**). Each colored scatter dot on top of each bar represents the average MEP value for one participant. All data is presented as the mean ± SEM.

### Corticospinal excitability during baseline

Before and after participants performed the experimental task, we assessed baseline (resting state) CSE. These results are shown in Figure 5. We used a two-by-two repeated measures ANOVA with factors MUSCLE (FDI and APB) and TIME (pre- and post-experiment). Briefly, the analysis did not indicate that CSE was differently modulated between the FDI (mean = 1.50 ± 0.24 mV) and APB (mean = 1.15 ± 0.22 mV) (main effect of MUSCLE; F_(1,15)_ = 1.00; p = 0.33). No main effect of TIME was found either (F_(1,15)_ = 2.17; p = 0.16) as CSE did not differ significantly between pre- (mean = 1.15 ± 0.15 mV) and post-experiment (mean = 1.49 ± 0.22 mV). Noteworthy, the analysis revealed significance for the interaction effect MUSCLE X TIME (F_(1,15)_ = 7.55; p = 0.015) although post-hoc exploration did not reveal significant differences between measurements (Figure 5). In sum, our results provide no evidence that baseline CSE was differently modulated before or after the experiment and between the FDI and APB muscles.

**Figure 6.**
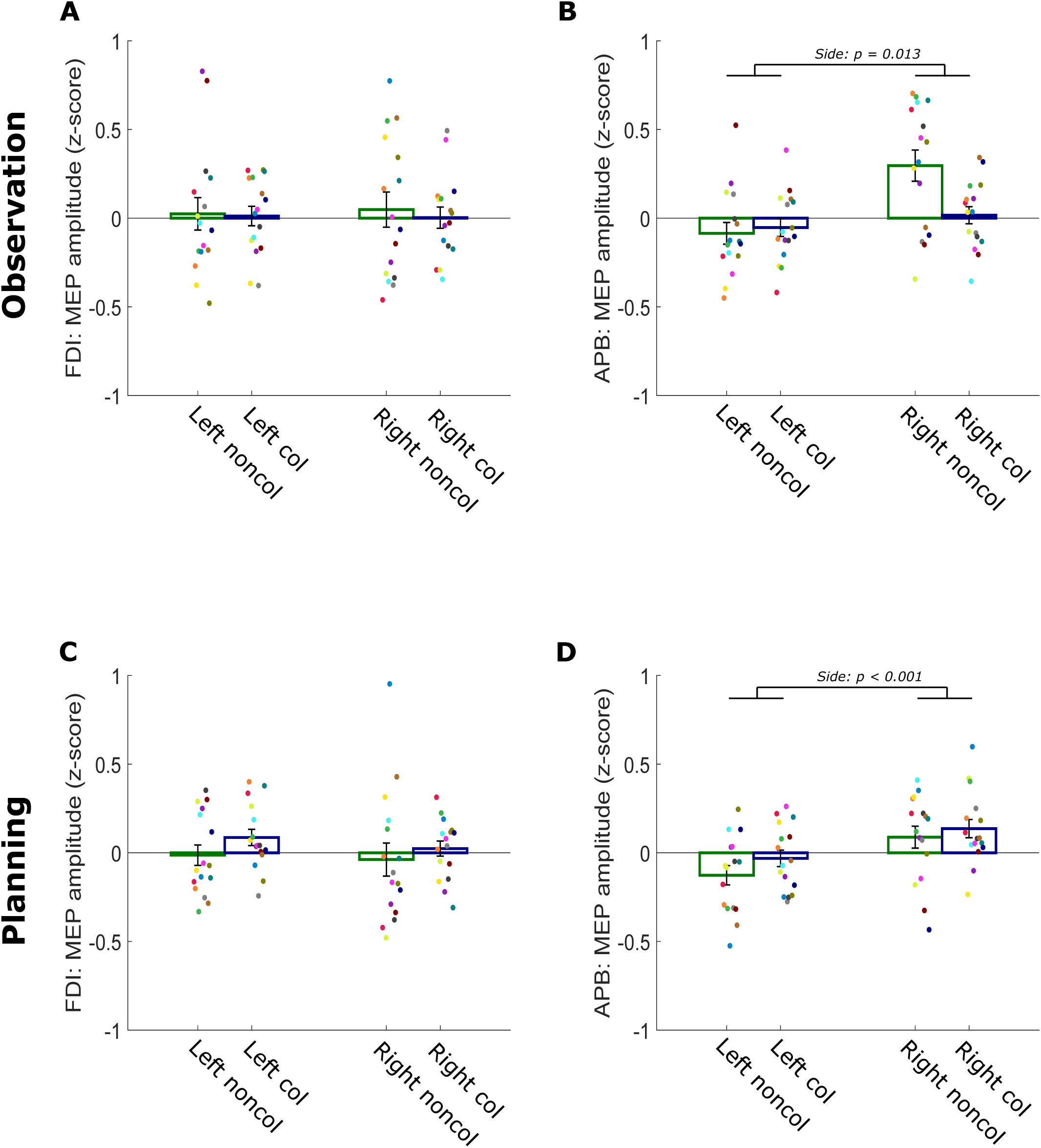
C**orticospinal excitability during lift observation and planning.** Average motor evoked potential (MEP) values (z-score normalized) during lift observation (**top row**) and lift planning, pooled for REPETITION, (**bottom row**) recorded from first dorsal interosseous muscle (**FDI; panels A and C**) and from the abductor pollicis brevis muscle (**APB; panels B and D**). The actor and participants used either collinear (col; blue) or noncollinear (noncol; green) digit positioning to lift the asymmetrical weight distributions (left or right). Each dot on top of each bar represents the average MEP value for one participant for that respective condition during lift observation and planning. All data is presented as the mean ± SEM.

### Corticospinal excitability during lift observation

As mentioned before, we hypothesized that modulation of motor resonance would be driven by specific observed movement features (i.e., digit positioning or muscle contraction).

#### First dorsal interosseus muscle (FDI)

As shown in Figure 6A, both the effect of SIDE and POSITION_SIDE_ were not significant (*both F* < 0.94, *both p* > 0.91). As such, our results provide no evidence that corticospinal excitability was differently modulated during the different observation conditions when being recorded from the FDI muscle.

#### Abductor pollicis brevis (APB)

In contrast to our findings for the FDI muscle, both the effects of SIDE and POSITION_SIDE_ were significant (*both F* > 5.19, *both p* < 0.008). As can be seen in Figure 6B, when participants observed the actor lifting the right-sided asymmetrical weight distribution, CSE was significantly larger (mean = 0.15 ± 0.06) compared to when participants observed lifts on the left-sided weight distribution (mean = -0.07 ± 0.04) (p = 0.013; *main effect of SIDE*). Although the nested effect POSITION_SIDE_ was significant, post-hoc comparisons did not reveal significant differences between conditions (*all p* > 0.06). In sum, these findings indicate that, CSE was significantly facilitated when observing lifts with an asymmetrical weight distribution that was right-sided.

### Corticospinal excitability during lift planning

Although this study was aimed to investigate motor resonance effects, we expected that CSE modulation would be similar (i.e., driven by the same movement features) as during lift observation.

#### First dorsal interosseus muscle (FDI)

All effects were not significant (*all F* < 1.42, *all p* > 0.24). As such, our results provide no evidence that CSE was differently modulated during the different lift planning conditions when being recorded from the FDI muscle (Figure 6C).

#### Abductor pollicis brevis (APB)

As shown in Figure 6D, in line with our findings for CSE modulation during lift observation, the effect of SIDE was significant (*F*_(1,192)_ = 16.23, *p* < 0.001) although all other effects were not (*all F <* 2.06, *all p* > 0.13). When participants planned to lift the right asymmetrical weight distribution (mean = 0.11 ± 0.04), CSE was significantly larger compared to when they planned to lift the left-sided one (mean = -0.08 ± 0.03) (p < 0.001; *main effect of SIDE*). As such, these findings indicate that, irrespective of digit positioning, CSE was significantly facilitated when planning to lift a right asymmetrical weight distribution.

### Background EMG during the experiment

To ensure that between-group and between-condition differences were not driven by differences in hand relaxation during lift observation and planning, we investigated potential differences in background EMG. For this we used the same statistics as the ones we used for lift observation and lift planning.

#### Background EMG during lift observation

For both the FDI and APB muscle, all effects were not significant (*all F* < 0.68, *all p* > 0.42). As such, these findings provide no evidence that background EMG different significantly between conditions when participants observed lifts on the asymmetrical weight distributions.

#### Background EMG during lift planning

For both the FDI and APB muscle, all effects were not significant (*all F* < 0.69, *all p* > 0.60). As such, these findings provide no evidence that background EMG different significantly between conditions when participants planned to lift the asymmetrical weight distributions.

## Discussion

In the present study, we investigated how CSE is modulated during observation of object lifting (i.e., ‘motor resonance’). Although we were initially interested in motor resonance effects only, we also recorded CSE during lift planning. Participants were asked to lift a manipulandum after observing an actor lift it. The object’s center of mass could be changed by placing a heavy cuboid in one of three compartments (Figure 1A). The object’s weight distribution could be ‘symmetrical’, by inserting the heavy cuboid in the middle compartment, or left- and right-sided ‘asymmetrical’, by inserting the heavy cuboid in the left or right compartment, respectively. Participants and actor were instructed to lift the object skillfully, i.e., generate appropriate compensatory torque to minimize object roll. Moreover, when lifting the asymmetrical weight distributions, digit positioning was constrained to two specific digit positioning strategies and, as mentioned before, TMS was applied during both observation and planning.

Our results indicate that the participants’ lift performance was suboptimal as they were not able to generate appropriate compensatory torque (Figure 4A). Noteworthy, the actor was also not able to do so when using collinear digit positioning on the asymmetrical weight distributions. As such, suboptimal lift performance was likely caused by limitations in our experimental set-up. Noteworthy, our results indicate that CSE modulation during both lift observation and planning was driven by the object’s asymmetrical weight distribution. That is, when participants observed or planned a lift on the right asymmetrical weight distribution, CSE recorded from the APB muscle was significantly increased compared to observing or planning a lift on the left asymmetrical one. During lift observation CSE modulation seemed to be primarily driven by an observed change in digit positioning (i.e., the thumb moving to the lower constrained location; Figure 4B). During lift planning, CSE modulation seemed to be more generally increased for both constrained digit positioning conditions on the right asymmetrical weight distribution (Figure 6D). In line with previous work (Alaerts, et al., 2010b; Rens et al., 2020), these findings indicate that motor resonance can be modulated by an object’s weight distribution similarly to other intrinsic object properties (such as weight).

Previous studies have shown that CSE is modulated during lift observation (‘motor resonance’). For instance, Alaerts et al. (2010a, 2010b) showed that motor resonance during lift observation reflects object weight as indicated by visual object properties (such as degree of filling) or by observed movement features (such as muscle contraction or movement kinematics). In addition, Buckingham et al. (2014) showed that motor resonance is driven by object size when observing skilled, but not erroneous lifts. Last, Rens et al. (2020), showed that motor resonance is modulated by object weight but can be strongly biased by contextual cues. Our findings corroborate these studies by showing that motor resonance was modulated by the intrinsic object properties. Specifically, CSE recorded from the APB muscle was increased when observing lifts in which the object’s weight was asymmetrically distributed to the right side (Figure 4). Considering that the object was visually identical in all weight distribution conditions, participants could not rely on visual object-related information (e.g., the heavy cube being visually different). As such, motor resonance should have been solely driven by observed movement features (such as digit positioning or object muscle contraction) within the observed lifts on the right asymmetrical weight distribution. However, it is important to note that CSE modulation was only present in the APB and not in the FDI muscle which might indicate that these effects of complex object properties (i.e., weight distribution) are relatively weak.

When the actor had lifted the symmetrical weight distribution and would then lift the right asymmetrical one with noncollinear positioning, she would shift her thumb on the light side to the lower constrained location whereas the index finger stayed on the same upper one. It is likely that this shift in digit positioning increased motor resonance when observing noncollinear lifts on the right asymmetrical weight distribution whereas the other movement features contributed arguably less. That is, when the actor lifted both asymmetrical weight distributions noncollinearly, she generated similar compensatory torques (Figure 3A). In addition, the actor used higher grip forces when lifting the asymmetrical weight distributions with collinear positioning (Figure 3C) and she scaled her thumb load forces higher when lifting the left-sided asymmetrical weight distribution with noncollinear positioning (Figure 3D). As such, only when observing noncollinear lifts on the right asymmetrical weight distribution did the thumb positioning change whereas the other movement features (i.e., force and compensatory torque) in this condition were similar to those in the noncollinear left asymmetrical condition. Given the change in digit positioning and the overlap in other movement features, it is likely that motor resonance effects in this condition were primarily driven by the observed digit positioning.

In contrast to our initial hypotheses motor resonance effects were statistically driven by differences in the asymmetrical weight distribution rather than constrained positioning conditions (Figure 6). Accordingly, motor resonance should also have been increased when observing lifts on the right asymmetrical weight distribution with collinear positioning. There are several factors that could have driven this increase in motor resonance. First, in this condition, the actor’s performance was the worst as she generated the least amount of compensatory torque (Figure 3A). Second, in this condition she scaled her grip forces higher than in all other conditions (Figure 3C). Third, the thumb had to generate almost no load force as this was done by the index finger on the heavy side (Figure 3D). Last, digit positioning did not deviate strongly from those in the other collinear conditions. As such, motor resonance effects when observing collinear lifts on the right asymmetrical weight distribution could have been driven by object roll (due to the inappropriate amount of compensatory torque) or the observed grip forces. Arguably, the contribution of the observed grip forces on motor resonance is debatable.

Specifically, when the actor lifted the left asymmetrical weight distribution with collinear positioning, she generated more load forces with her thumb compared to when she used the same positioning for the right asymmetrical weight distribution (Figure 3D). As such, this ‘force interpretation’ would suggest that observed grip forces modulate motor resonance whereas observed load forces do not. Although Alaerts et al., (2010b) have demonstrated that CSE is modulated during observed squeezing (i.e., visible muscle contraction: grip forces) and during observed lifting when the hand contraction cannot be seen (i.e., movement kinematics: indicative of the load forces), it is still unsure how these force parameters converge in modulating CSE during observed lifting. To our knowledge, the differential muscle contractions of grip and load forces on CSE modulation have not been disentangled yet.

To end, we initially hypothesized that motor resonance would be selectively modulated by either observed digit positioning or muscle contractions. However, our results suggest that modulation of motor resonance was more generally modulated by movement features indicating the object’s right asymmetrical weight distribution. Indeed, motor resonance was primarily increased when observing a visible shift in digit positioning in the right noncollinear condition and by the largest object roll, caused by inappropriate compensatory torque, in the right collinear condition.

Based on previous findings of our group (Rens et al., 2020), it is plausible that digit positioning and object roll could have modulated motor resonance. In that study, we highlighted that motor resonance is only modulated by object weight when the actual object weight would always match the participants’ weight expectations. However, when the participants’ weight expectations could be incorrect, motor resonance was rather driven by a mechanism monitoring these expectations. Those findings have been supported by a reasoning within the review of Amoruso and Finisguerra (2019). They argued that motor resonance reflects the inner replica of the observed action when observed in isolation but can be altered by higher-level factors (such as contextual cues) when present. In sum, both these works indicate that motor resonance can be flexibly driven by different movement features during action observation. In the current study, participants were asked to minimize object roll during lift observation. This contextual importance of accurately estimating the weight distribution during lift observation might have caused motor resonance to be driven by ‘salient’ movement features indicating the object’s weight distribution. Arguably, it is plausible that participants focused on shifts in digit positioning and visible object roll to perceive the weight distribution which in turn modulated their motor resonance.

Critically, we did not find similar modulation of CSE recorded from the FDI muscle. This was likely caused by experimental limitations. When the inverted T-shape manipulandum was placed in front of the participant, it was rotated (< 45 degrees) according to the participants’ preferences. As most participants found the manipulandum relatively heavy, the rotated position enabled them to reduce wrist overextension, improving lifting comfort. Due to the object rotation, the index finger was hidden behind the manipulandum and participants had no vision on the index finger during both lift observation and execution. These visiblity limations might have completely eradicated any CSE modulation recorded from the FDI muscle. It is also possible that similar visiblity limitations drove the selective modulation of motor resonance for the right but not left asymmetrical weight distribution. When the actor lifted the manipulandum (see dyadic positioning; Figure 1C), she would reach with her arm in front of the participant blocking vision on the manipulandums left side with her lower arm. Because of these visiblity limitations, participants might have primarily focused on thumb actions and the right side of the manipulandum. However, it is important to note that the participant’ lift performance for the noncollinear conditions did not differ between asymmetrical weight distributions. Furthermore, when using collinear positioning, participants performed worse on the right asymmetrcial weight distribution compared to the left one. As such, even though motor resonance was selectively modulated for the right asymmetrical weight distribution, participants perceived both weight distributions similarly during lift observation as indicated by their own lift performance after lift observation.

In a recent motor resonance study on humans, Cretu et al. (2019) investigated whether motor resonance effects are present when the hand-object interaction cannot be seen. They found that motor resonance can be driven by contextual cues but only if they are informative of the hidden action. As such, Cretu et al. (2019) showed that that relevant information regarding the observed action needs to be present in order to modulate motor resonance. In addition, we previously showed that motor resonance is flexibly modulated based on experimental context (Rens et al., 2020). As participants in the present study had no vision on the index finger during observed object lifting, they may have focused solely on the thumb thus eradicating motor resonance effects recorded from the FDI muscle. However, future research is necessary to substantiate the notion that motor resonance can be digit-specifically modulated based on visibility. Additionally, futures studies could place the actor opposite of the participant or use transparent objects to ensure both fingers are always visible. We initially decided to place the actor at the side of the participant as Alaerts et al. (2009) showed that motor resonance effects are stronger when executed actions are observed from a first person point of view.

With respect to lift execution, our participants were not able to lift the manipulandum skilfully in the collinear condition, even after haptic feedback (i.e., second and third lifts; Figure 4A). In contrast, in the noncollinear condition, participants were able to generate appropriate compensatory torque already in their first lift (Figure 4A) and did not improve from their first to second lifts. Critically, our actor also performed suboptimally when lifting collinearly. In contrast, in Fu et al. (2010), participants were able to lift an asymmetrical weight distribution skilfully when using constrained collinear digit positioning. Taken together, it may not have been feasible in our study to lift the asymmetrical weight distribution skilfully with collinear digit positioning. It is important to note that even though our manipulandum and the one of Fu et al. (2010) had similar external torque, ours was slightly heavier. In addition, it is possible that textural differences (i.e., friction of the graspable surfaces) differed between ours and theirs which led to differences in performance (Johansson & Westling, 1984). As such, having decreased the object weight, thus making lifts on the asymmetrical weight distributions less challenging, could have removed the potential confounding effects of lifting performance differences between conditions.

Parikh et al. (2014) showed that when individuals plan to generate high or low forces, CSE during planning is decreased when planning to generate high forces. Their findings suggest that predictive force planning is force-dependently modulated akin to work of Loh et al. (2010) on predictive object lifting. However, in our study this force-dependent modulation seemed to be absent: when participants planned to lift the right asymmetrical weight distribution, CSE was similarly modulated in both digit positioning conditions even though they generated more force in the collinear than in the noncollinear condition. As such, CSE during lift planning was unlikely modulated by planned forces but rather by the object’s weight distribution. However, as lift planning was contaminated by suboptimal performance, future research is necessary to substantiate whether the motor system can veridically encode an object’s weight distribution during lift planning. Furthermore, Davare et al. (2019) showed that CSE is increased when lifting an asymmetrical weight distribution with unconstrained compared to constrained digit positioning. This suggests that CSE modulation is driven by the sensorimotor uncertainty (regarding digit positioning). However, this effect was only present when TMS was applied during early object contact and not during mid-reach. With respect to our study, it is unlikely that mechanisms related to sensorimotor uncertainty affected CSE modulation during lift planning as we constrained digit positioning in all conditions and we applied TMS during lift planning, not early object contact.

In our study, we decided on our constrained digit positionings with the intention of having the digit positioning difference and load force difference approximate zero in the collinear and noncollinear conditions, respectively (in support of this statement see Figure 3 for the actor’s performance). However, as mentioned before, lifting performance was suboptimal for both actor and participants. As such, future studies could decrease object weight or let participants choose their own constrained digit positionings to ensure skilled performance. In our previous work (Rens et al., 2021), we found that observing skilled lifts of asymmetrical weight distributions did not enhance predictive lift planning in the observer. In the present study, we found that participants generated more compensatory torque after observing skilled lifts with noncollinear digit positioning, contrasting the findings of our previous study. Arguably, this difference is driven by the constrained digit positioning in the present study as participants were free to choose their own digit positioning in Rens et al. (2021). However, more research is required to investigate the interaction between action observation and constraining motor execution on predictive lift planning.

Even though lifting performance was suboptimal, our results indicate that CSE modulation during lift planning and observation is not associated with the participants’ lift performance. Although CSE was similarly modulated for collinear and non-collinear lifts on the right center of mass, participants did not lift the right asymmetrical weight distribution skillfully with collinear digit positioning. Indeed, a limitation in our study is the erroneous lifting performance on the asymmetrical weight distribution when using collinear digit positioning: Buckingham et al. (2014) showed that motor resonance can be differently modulated when observing skilled or erroneous actions. As such, future research is required to disentangle how motor resonance relates to motor planning and proper execution.

Last, it has been shown that S1 is involved in integrating haptic feedback to generate appropriate load forces when lifting objects (Parikh et al., 2020) and stimulating it during our task could have affected lift performance. The scalp location for stimulating S1 has been shown to be located approximately 2 cm lateral and 0.5 cm posterior to the scalp location for stimulating M1 (Holmes et al., 2019; Holmes & Tamè, 2019). However, confounding stimulation of S1 may be unlikely as we oriented the TMS coil to induce a posterior-anterior current. In addition, the spread of TMS across neighboring tissue has been considered relatively small based on modelling (Deng et al., 2013) and single-cell recordings in the macaque monkey (Romero et al., 2019). In particular, Romero et al., (2019) showed that the spread of single-pulse TMS was limited to less than 2 mm in diameter. Although we cannot exclude that S1 was unintentionally stimulated in the present study, it should not have affected conditions differently. That is, we applied TMS during every trial (i.e., during lift observation and during lift planning on all weight distributions). As such, potential confounding effects of S1 stimulation should have affected all conditions equally. To end, to investigate this potential confounding effect of TMS, future studies could include a ‘no stimulation condition’ to compare lift performance with or without single-pulse TMS during lift planning.

Motor resonance has been argued to rely on mirror neurons. Mirror neurons are similarly activated when executing or observing the same action and have been argued to be involved in action understanding by “mapping” observed actions onto the cortical representations involved in their execution (Rizzolatti et al., 2014). Mirror neurons are primarily located in M1, the ventral premotor cortex (PMv) and the anterior intraparietal area (AIP) (Rizzolatti et al., 2014). Importantly, these regions also constitute the cortical grasping network which is pivotal in planning and executing grasping actions (for a review see Davare et al. 2011) further substantiating these neurons involvement in action understanding. Our findings corroborate mirror neuron functioning by showing that CSE was similarly modulated during lift observation and planning. Interestingly, even though our motor resonance findings might have been partially driven by the experimental context (Rens et al., 2020), CSE modulation was similar during planning, which substantiates that the same mechanisms underlied CSE modulation in our experiment. To end, CSE modulation being similarly increased for the right asymmetrical weight distribution during both observation and planning indicates that the motor system encoded the same object-related information during both planning and observation.

In conclusion, the present study investigated how CSE is modulated during the observation and planning of lifts with an asymmetrical weight distribution. Our findings suggest that CSE modulation during observation was driven by the object’s weight distribution as potentially indicated by digit positioning and object roll (i.e., inappropriate compensatory torque). During lift planning, CSE was similarly modulated by the object’s weight distribution indicating, in line with previous research (for a review see Cattaneo & Rizzolatti, 2009), that the same neural mechansisms are involved in CSE modulation during lift observation and planning. As such, our findings provide further support that the motor system is involved in the observation and planning of hand-object interactions.

## Acknowledgements

We are grateful to Dr. Massimo Penta (Arsalis, Belgium) for the design of the manipulandum and Isa Vanstraelen for her help in data collection. This work was funded by a Research Foundation Flanders (FWO) Odysseus Project (Fonds Wetenschappelijk Onderzoek, Belgium: G/0C51/13N) awarded to MD and 12X7118N/Research Foundation Flanders (FWO) awarded to VVP.

## Notes

### Competing Interest Statement

The authors have declared no competing interest.

